# Functional rescue of F508del-CFTR through revertant mutations introduced by CRISPR base editing

**DOI:** 10.1101/2024.08.28.610115

**Authors:** Irene Carrozzo, Giulia Maule, Carmelo Gentile, Alessandro Umbach, Matteo Ciciani, Daniela Guidone, Martina De Santis, Gianluca Petris, Luis Juan Vicente Galietta, Daniele Arosio, Anna Cereseto

## Abstract

Cystic Fibrosis (CF) is a life-shortening autosomal recessive disease caused by mutations in the *CFTR* gene, resulting in functional impairment of the encoded ion channel. F508del mutation, a trinucleotide deletion, is the most frequent cause of CF affecting approximately 80% of patients. Even though current pharmacological treatments alleviate the F508del-CF disease symptoms there is no definitive cure. Here we leveraged revertant mutations (RMs) in *cis* with F508del to rescue CFTR protein folding and restore its function. We developed CRISPR base editing strategies to efficiently and precisely introduce the desired mutations in the F508del locus. Both editing and CFTR function recovery were verified in CF cellular models including primary epithelial cells derived from CF patients. The efficacy of the CFTR recovery strategy was validated in cultures of pseudostratified epithelia from patients’ cells showing full recovery of ion transport. Additionally, we observed an additive effect by combining our strategy with small molecules that enhance F508del activity, thus paving the way to combinatorial therapies.

## Introduction

Cystic Fibrosis (CF) is a life-shortening autosomal recessive disease that affects at least 100.000 people worldwide^1,2^. It is caused by mutations in the cystic fibrosis transmembrane conductance regulator (*CFTR*) gene, which encodes a transmembrane channel localized at the apical surface of the epithelial cells^3^. CFTR is fundamental to keep correct homeostasis across epithelial membranes by assuring mucus hydration and clearance in particular in the airways. Indeed, CFTR impairment affects multiple organs, yet morbidity and mortality of CF patients are mainly caused by progressive obstructive lung disease^1,2^.

According to the CFTR2 database^4^, more than 700 mutations in the *CFTR* gene are causative of CF. The most common mutation is the F508del (present in 80% of CF patients), a deletion of 3 nucleotides that determines the loss of a phenylalanine residue^2,4^. This genetic alteration results in a misfolded CFTR protein that is targeted for degradation by the ubiquitin-proteasome system^5^. A small fraction of mutated CFTR can evade early degradation and reach the plasma membrane (PM), but it remains insufficiently active and unstable. Efficient CF therapies were developed consisting of small molecules that modulate CFTR folding (correctors) or gating activity (potentiators)^2^. The current recommended treatment for patients carrying the F508del mutation (in one or both alleles) is a combination of three modulators elexacaftor-tezacaftor-ivacaftor (ETI) consisting of two correctors and a potentiator^2^. Although ETI significantly improves patients’ health, it is not a definitive cure and requires lifelong administration. As often occurring with pharmacological regimens, these treatments can cause side effects and occasionally may not produce the desired therapeutic efficacy due to various factors, including genetic variability^6,7^

Genome editing is a new powerful instrument for gene therapy to treat genetic diseases^8^. In CF various genome editing strategies have been developed to repair CFTR mutations including the F508del defect^9^. Given the nature of this mutation consisting in a small deletion, several groups used the homology directed repair (HDR) strategy consenting the replacement of sequences favored by programmable nucleases, including zinc finger nucleases or Cas9^10–15^. Yet this method is limited to specific target cells as it depends on specific phases of the cell cycle (S-G2/M) and is associated with risks of genotoxicity induced by the DNA double strand breaks (DSB)^16–18^. Recent technological advancements in genome editing include the development of prime editing^19^, a DSB-free approach that was efficiently used to correct F508del reaching up to 50% function compared to wild-type^20,21^. Nonetheless, prime editing generates a high percentage of unwanted small deletions/insertions (indels) with unpredicted outcomes^20,21^ and its composition with multiple elements increases the molecular size of the delivery cargo thus challenging the *in vivo* treatments^22,23^.

Previous studies showed that revertant mutations (RMs) in *cis* with F508del can stimulate proper folding and function of CFTR protein^24^. The first RM was identified in 1991 with a patient reporting a milder form of CF^25^. The genetic analysis revealed a compound F508del-R553Q and R553X, the latter corresponding to a nonsense mutation. Therefore, the milder phenotype was very likely generated by the second-site mutation in the F508del-*CFTR* locus which at least partially abrogated the CF phenotype. This observation prompted the search for additional RMs leading to the identification of I539T, G550E, R555K and R1070W^26–32^. Most of these RMs are located within the nucleotide binding domain (NBD1) of CFTR, except for the R1070W, which resides in the fourth intracellular loop (ICL4) (**Figure 1A**). RMs in the NBD1 reduce the thermodynamic and kinetic instability during F508del-CFTR biogenesis^29–31^, while R1070W restore the ICL4/NBD1 interface^30,32^.

**Figure 1.**
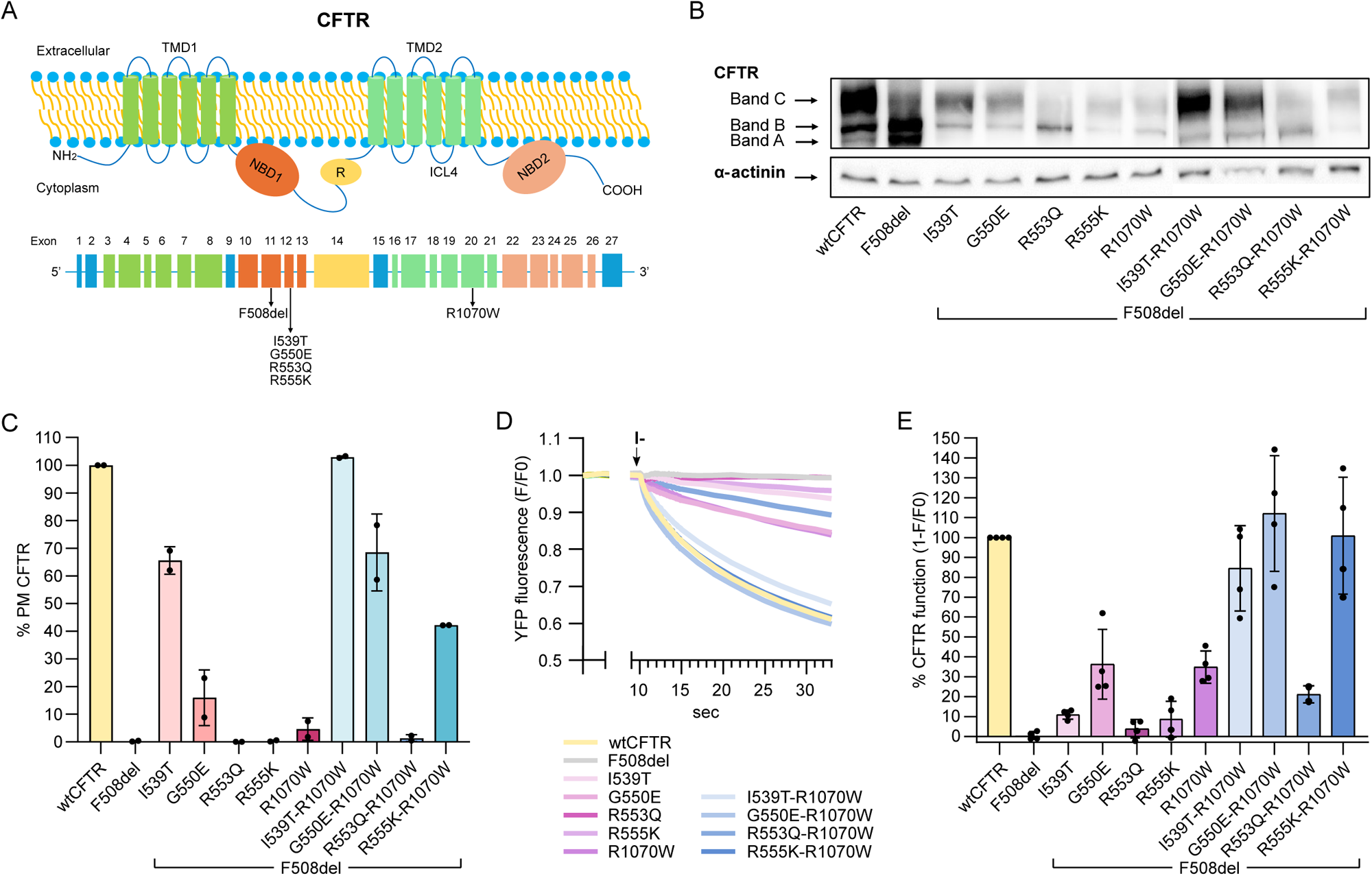
RMs rescue F508del-CFTR maturation, localization and function. (A) Schematic representation of CFTR protein structure, highlighting its five domains: two transmembrane domains (TMD1 and TMD2), each formed by six transmembrane segments, two cytosolic nucleotide binding domains (NBD1 and NBD2) and the regulatory domain (R). The lower panel shows exons and the locations of the F508del and RMs within the CFTR locus. Corresponding protein domains are color coded. (B) Representative western blot analysis of CFTR produced by the F508del modified with the indicated RMs and transduced in HEK293T cells with lentiviral vectors. The blot was developed with anti-CFTR antibodies, upper panel, or L-actinin, lower panel. Bands A, B and C are indicated by the arrows. (C) Histograms representing the percentage of CFTR (3HA-tagged) rescued at the PM detected by flow cytometry. Samples are normalized to wtCFTR, which is considered 100% at the PM. Data are means ± SD from n=2 independent experiments. Gating strategy is shown in **Figure S1A**. (D and E) HS-YFP quenching assay performed in HEK293T stably expressing HS-YFP and transfected with the indicated CFTR variants. (D) Representative traces of HS-YFP assay recorded with a fluorescence microscope (see Methods). Cells were stimulated with 10 µM forskolin before the addition of iodide (I-), indicated by the arrow. (E) Quantification of CFTR activity as a measure of HS-YFP quenching from n=4 independent experiments. CFTR function is calculated as 1-F/F0 using values recorded 7 seconds after I-addition. Data are means ± SD.

Here we exploit CRISPR base editors to introduce RMs into the F508del-*CFTR* locus as a strategy to restore CFTR activity. The efficiency, precision and competitive molecular size among the genome editing tools offer the opportunity for a definitive cure to F508del CF patients.

## Results

### Restoration of F508del defect mediated by revertant mutations in CF cell lines

We evaluated the impact of RMs I539T, G550E, R553Q, R555K and R1070W on F508del-CFTR by first analyzing the protein maturation. Different CFTR maturation products are generated during the protein channel biogenesis which can be detected through western blot analysis: a non glycosylated form, band A (135 kDa), an immature partly glycosylated form, band B (150 kDa), and a fully glycosylated mature form, band C (170 kDa)^33^. The F508del-*CFTR* carrying the RMs, single or in combinations, were expressed in HEK293T cells using lentiviral vectors. The I539T-R1070W and G550E-R1070W induced the formation of mature CFTR proteins at levels comparable to wild type CFTR, while partial maturation rescue was observed with the single I539T, G550E, R555K, R1070W, and combined R555K-R1070W RMs (**Figure 1B**).

To further evaluate the rescue efficacy of the RMs on F508del-*CFTR,* we analyzed CFTR cellular localization, which is physiologically at the plasma membrane (PM) if not impaired by mutations that alter protein folding^34^. We performed flow cytometry analyses using a 3HA tag located at the fourth external loop of CFTR. The amount of CFTR restored at the PM (**Figure 1C** and **S1A**) mirrored the levels of maturation observed by western blot, showing a strong localization rescue by the combinations I539T-R1070W and G550E-R1070W and also by the I539T while the remaining mutations showed variably lower levels of rescue.

Finally, to assess the CFTR function we performed the halide-sensitive YFP (HS-YFP) assay^35,36^ in HEK293T cells. This assay exploits the quenching property of HS-YFP by iodide anions. In cells where CFTR is functionally active, iodide ions enter the cells through the CFTR channels, leading to the quenching of HS-YFP which results in fluorescence decrease, while in CFTR negative cells the fluorescence remains unaltered. This assay was performed in HEK293T stably expressing HS-YFP and transfected with various F508del-CFTR variants. Consistent with the maturation and cellular localization data, the I539T, G550E, R555K combined with R1070W rescued CFTR activity to near wild type levels. Partial or no rescue was observed with the single revertants, and the combination R553Q-R1070W (**Figure 1D**).

### CRISPR-base editing strategies to introduce revertant mutations in F508del-*CFTR*

CRISPR-base editor strategies were developed to introduce RMs I539T, G550E, R553Q, R555K and R1070W in the F508del-*CFTR* locus (**Figure S1B**). We used HEK293T cells carrying the F508del-*CFTR* cDNA to monitor the incorporation of specific RMs and to assess any nucleotide modifications occurring beyond the primary target, known as bystander edits^37^.

Depending on the RM to be inserted, we used either adenine base editors (ABE)^38^ or cytosine base editors (CBE)^39^ for A>G or C>T substitutions, respectively, utilizing different versions of nucleotide deaminases^40–42^. By leveraging the highest compatibility with protospacer adjacent motives (PAM) recognition sequences surrounding the target sites, we tested base editors made with three SpCas9 variants, SpCas9-NG^43^, SpG or SpRY^44^. For each base editor several sgRNAs were tested, indicated with numbers corresponding to the position of the target nucleotide relative to the PAM sequence^45^.

To insert the I539T mutation we tested several ABE versions (ABE8e, ABE8.20m and ABEmax)^40–42^. We obtained variable levels of A>G conversions at the target nucleotide, with up to 45% of editing with SpG-ABEmax, SpG-ABE8.20m and SpG-ABE8e (**Figure 2A**). Most of the ABEs tested showed substantial bystander activity in position A4, and A9, except for NG-ABEmax and SpG-ABEmax, which showed minimal conversion at A4 and less than 10% conversion at A9, resulting in a silent N537N mutation.

**Figure 2.**
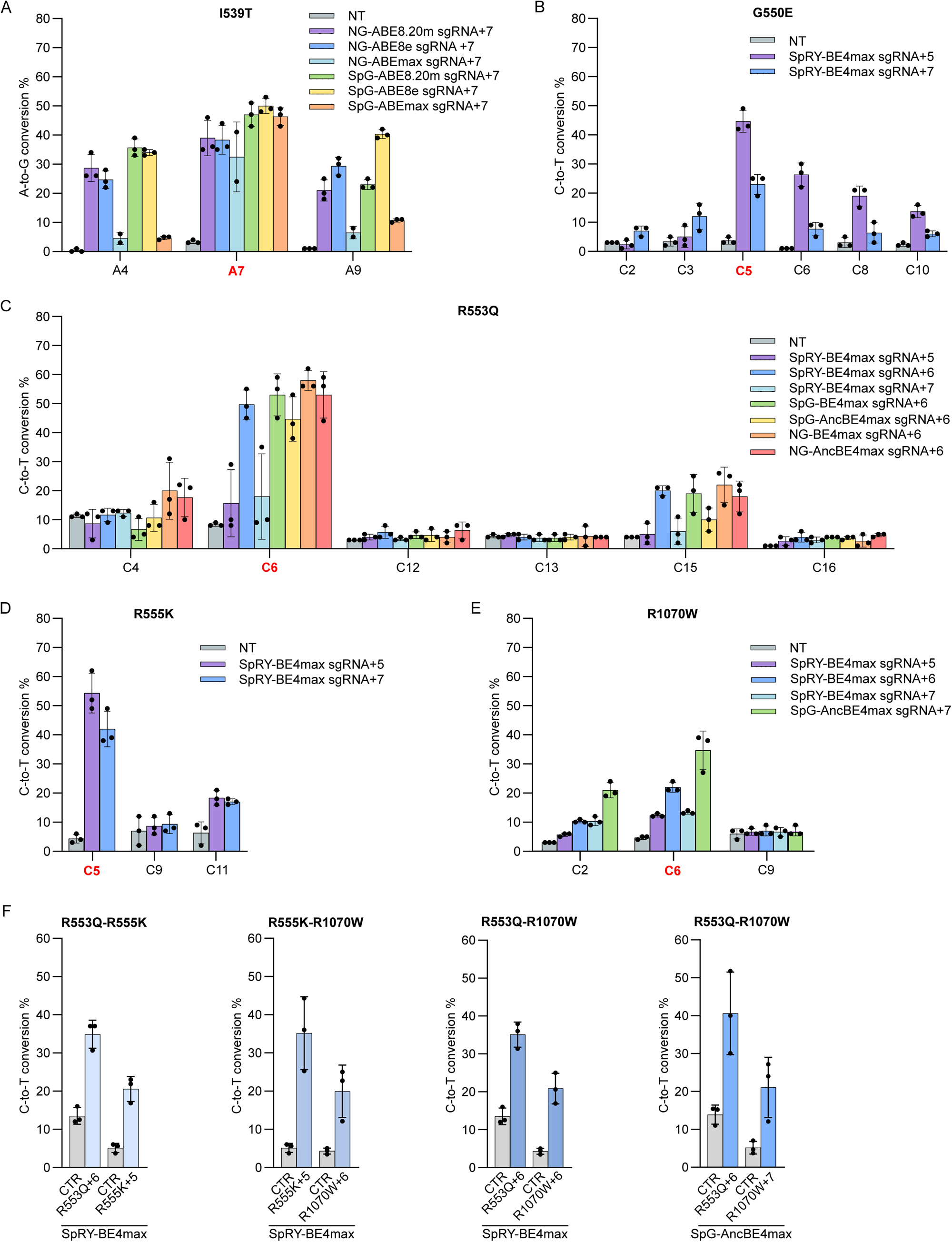
Insertion of RMs in the F508del-*CFTR* locus through CRISPR base editors. (A-F) Editing efficiency in F508del-*CFTR* HEK293T cells transfected with the indicated base editors and sgRNAs used (A-E) individually or (F) in combination to generate RMs indicated at each graph. Editing was quantified by EditR tool^43^ after PCR amplification and Sanger sequencing of the target locus. The target base for each RM is highlighted in red; bystander edits are shown in black. Data are means ± SD from n=3 independent experiments.

To insert G550E, R553Q, R555K and R1070W mutations we used two CBE versions, BE4max^41^ and AncBEmax^41^, combined with different sgRNAs and SpCas9 variants (**Figure 2B-E**).

For the G550E mutation, we obtained up to 45% editing at the target cytosine (C5), however associated with relevant bystander edits throughout the base editing window (**Figure 2B**). Two specific bystander edits, C2 and C8, are predicted to have a major functional impact since they produce CF-causing mutations (G551D and S549N)^46^ (**Figure 2B**).

The R553Q was introduced with up to 50% efficiency at C6 using various combinations of CBEs and sgRNAs (**Figure 2C**) showing main bystander edits at C15 (**Figure 2C**). Interestingly, the C15 bystander introduces another RM, the G550E, which is beneficial to rescue the F508del-*CFTR* folding. The most efficient editing strategies, as a result of both specific and synergistic bystander editing, were obtained with SpRY-BE4max and SpG-BE4max with sgRNA+6, generating simultaneous introduction of G550E-R553Q RMs.

The R555K was generated with SpRY-BE4max in combination with two different sgRNAs. Target base conversion reached over 50% efficiency with sgRNA+5 (**Figure 2D**), with the advantage of introducing a second RM, the R553Q through a bystander edit in C11, thus resulting in double R553Q-R555K RMs. The R1070W mutation was introduced using SpRY-BE4max and SpG-AncBE4max, resulting in up to 30% editing at C5. Additionally, a bystander edit at C2 generated a silent mutation, the F1068F (**Figure 2E**).

Next, we explored the possibility of using a single base editor to introduce combinations of RMs. The possible compatible combinations were R553Q-R555K, R553Q-R1070W and R555K-R1070W, while I539T and G550E could not be inserted together with others. The R553Q-R555K, R553Q-R1070W and R555K-R1070W were inserted using SpRY-BE4max, while the R553Q-R1070W with SpG-AncBE4max. The RM combinations were overall efficiently edited (**Figure 2F**). We observed a more evident reduction of R555K editing when combined with the R553Q, probably due to sgRNAs interference (**Figure 2F**). Bystander edits were similar to single guide strategies (**Figure S2**).

### Rescue of F508del-CFTR with revertant mutations introduced via base editing

We evaluated the rescue of the F508del-CFTR modified by using the most efficient base editing strategy identified in **Figure 2**. First, we analyzed PM localization which showed partial rescue (5%-31% compared to wild type) with all RMs (**Figure 3A**). Interestingly, the individual R553Q and R555K showed higher PM recovery than the corresponding modifications in F508del-*CFTR* cDNA (compare with **Figure 1C**). The higher PM recovery was probably due to bystander editing introduced by the additional RMs as reported in **Figure 2C-D**. We also assessed PM localization with combinations of RMs which showed higher CFTR rescue, with the maximum generated by R555K-R1070W reaching 40% of the wild type level (**Figure 3B**). The R553Q-R555K RMs showed decreased PM localization than R555K introduced individually (**Figure 3A** and **B**), probably due to interference between the sgRNAs (see **Figure 2D** and **F**).

**Figure 3.**
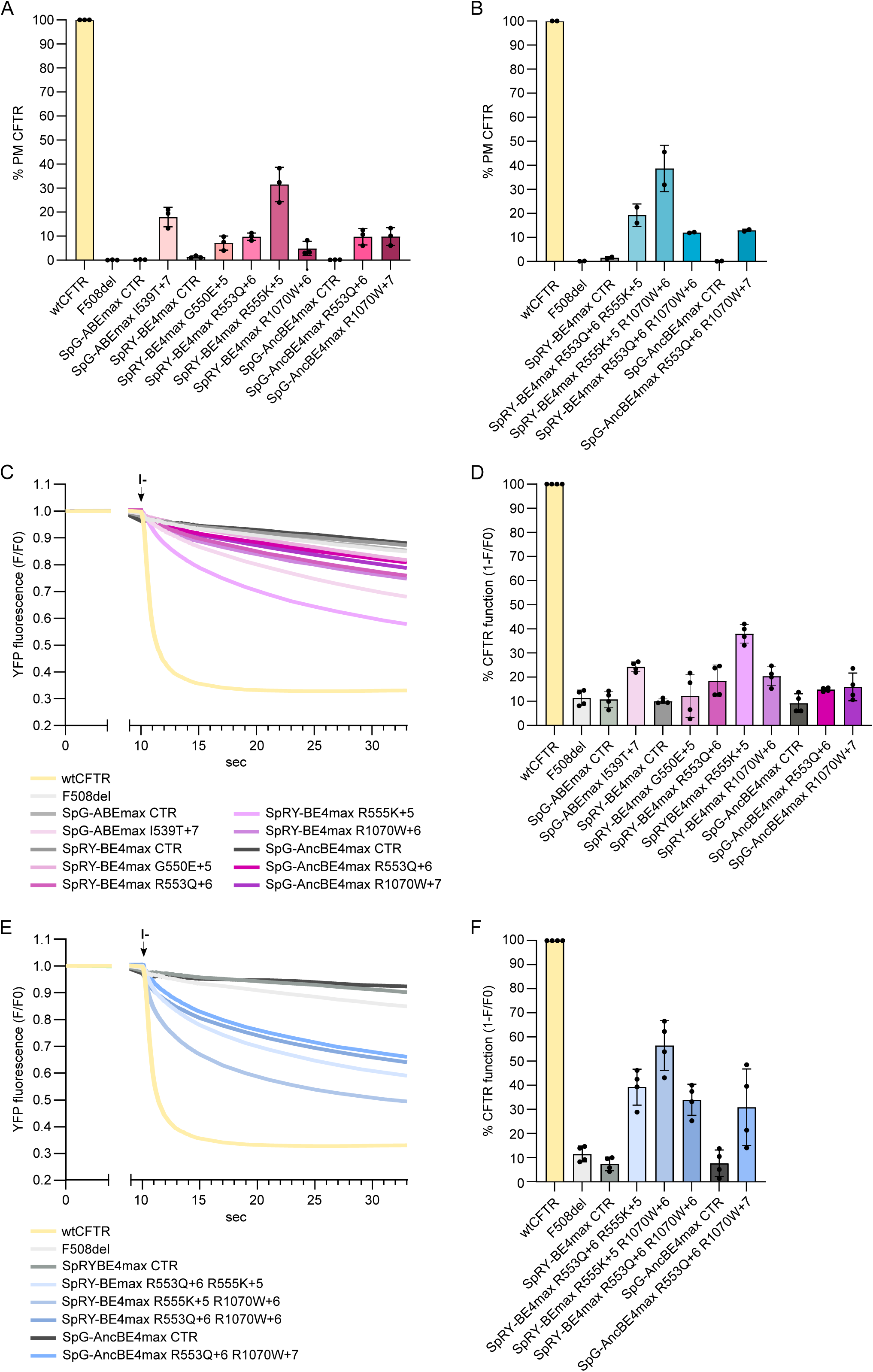
Rescue of F508del-CFTR localization and function induced by insertion of RMs. (A and B) Percentage of CFTR at the PM measured by flow cytometry in wtCFTR and F508del-*CFTR* HEK293T cells modified with the indicated base editors producing (A) single or (B) double RMs. Data are means ± SD from n=3 independent experiments. Samples are normalized to wtCFTR, which was set at 100%. (C-F) HS-YFP quenching assay in HEK293T endogenously expressing HS-YFP and F508del-CFTR cDNA and transfected with the indicated base editors and sgRNAs to insert (C,D) single or (E,F) double RMs. Cells were stimulated with 10 µM forskolin before addition of iodide, indicated by the black arrow. (C,E) Representative traces of HS-YFP quenching assay recorded through fluorescence microscopy. (D,F) Quantification of CFTR function as 1-F/F0 obtained 7 seconds from iodide addition. Data are means ± SD from n=4 independent experiments.

CFTR function was then evaluated through HS-YFP quenching assay. Among single RMs, R555K showed the highest activity (up to 40%) (**Figure 3C, D**), while with the double RMs, R555K-R1070W showed up to 56% of wild type activity (**Figure 3E, F**), paralleling the results observed with PM analysis.

### Combinatorial rescue of F508del-CFTR with RMs and CFTR modulators

We investigated the potential synergism between RMs and pharmacological treatments, including the corrector lumacaftor, which enhances CFTR folding, and the potentiator ivacaftor, which improves CFTR gating. To this aim, we performed the HS-YFP quenching assay in HEK293T cells stably expressing the variants of F508del-CFTR cDNA carrying the RMs.

We observed that individual RMs enhanced the activity of F508del-CFTR when treated with either modulators alone or in combination (**Figure 4** and **S3**), with the exception of R553Q, where no synergism was observed. For the majority of combined RMs, I539T-R1070W, G550E-R1070W and R555K-R1070W, which already increased F508del-CFTR activity close to wild type levels, we detected only a modest improvement of CFTR function when modulators were added. Conversely, RM combinations with lower baseline activity, R553Q-R555K and R553Q-R1070W, synergized with both modulators, reaching wild type levels of activity.

**Figure 4.**
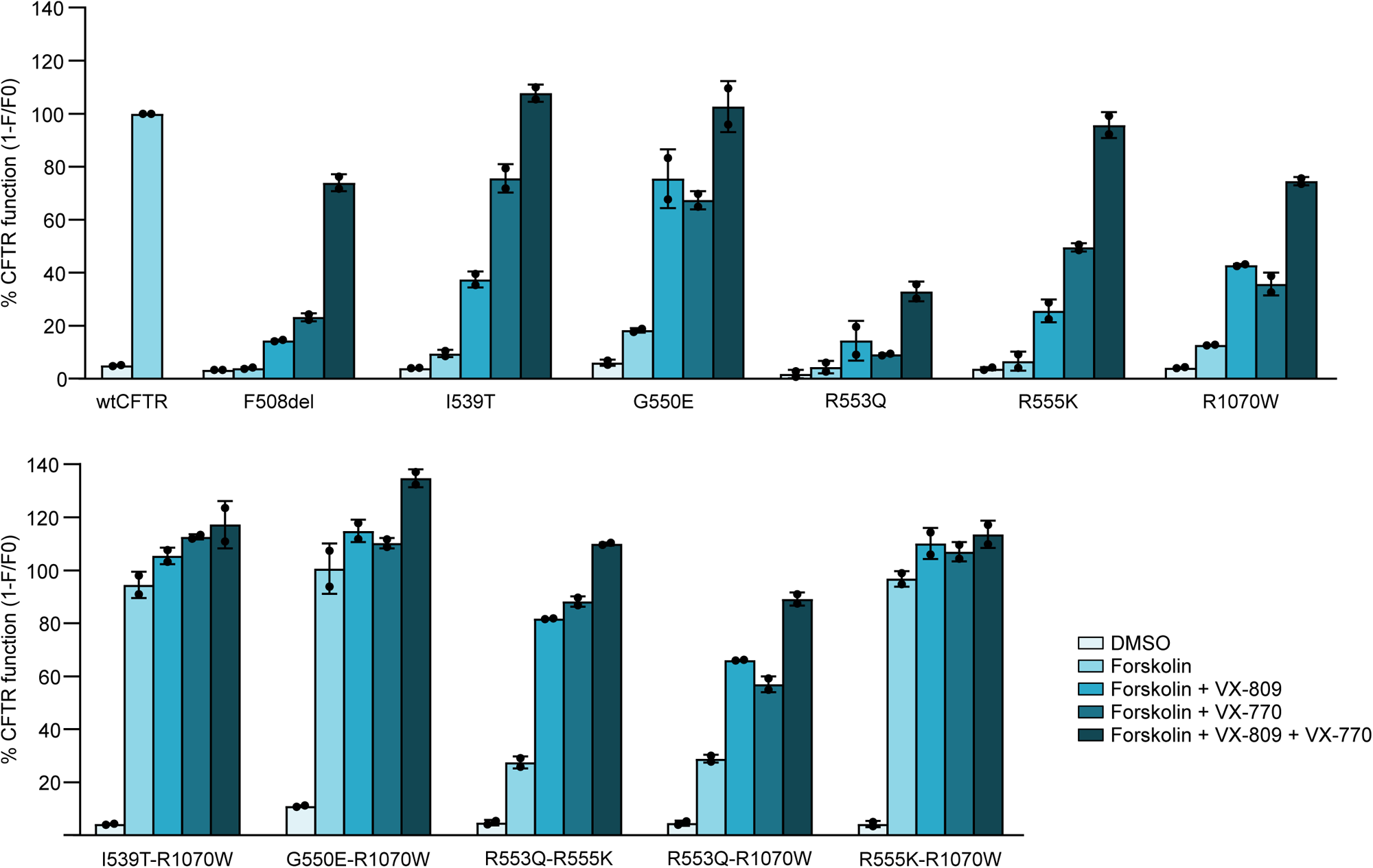
Evaluation of synergistic effects of RMs and modulators by HS-YFP quenching assay. Quantification of CFTR activity through HS-YFP quenching assay performed in HEK293T cells stably expressing F508del-CFTR variants and transfected with HS-YFP. Cells were treated either with forskolin, lumacaftor and ivacaftor, individually or in combination. CFTR function is expressed as 1-F/F0 based on fluorescence values recorded 7 seconds after iodide addition. Data are means ± SD from n=2 independent experiments.

### Efficiency and precision of F508del-*CFTR* base editing in patients derived primary bronchial epithelial cells

The base editing strategy was tested in CF models, consisting in culture of primary bronchial epithelial cells (HBE) derived from a patient homozygous for the F508del mutation.

Base editors were delivered by electroporation of RNA for both mRNA and sgRNA modified for optimal efficacy (see Methods) and by using specific ratios (**Figure S4A**, **B**). The efficacy of F508del modification with single or double RMs was as efficient as in the HEK293T models (**Figure 5A-E**). RNA delivery of base editors reduces bystander activity as formerly reported^45^, likely due to transient expression. This reduction was not observed for the I539T mutation, which showed off-target editing at A9, nevertheless leading to the generation of a silent mutation (**Figure 5A**), as observed in HEK293T cells (**Figure 2A**).

**Figure 5.**
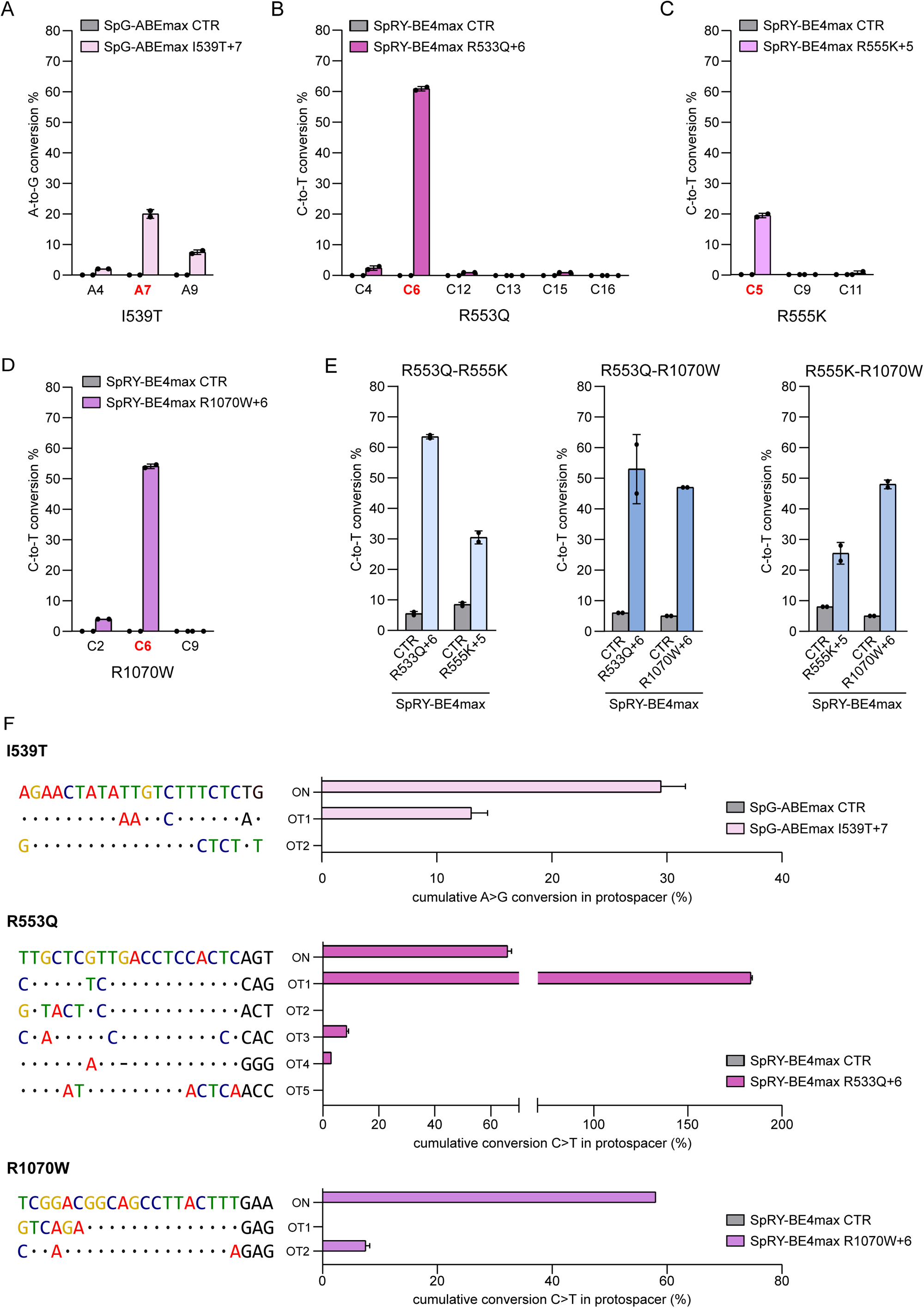
Insertion of RMs in HBE cells using base editors and evaluation of editing precision. (A-D) Editing efficiency for the insertion of single RMs in HBE cells homozygous for the F508del-CFTR gene. Editing was quantified by deep sequencing analyses. HBE were electroporated with mRNA encoding the indicated base editors and chemical modified sgRNAs specific for the target nucleotide. The target nucleotides are highlighted in red; bystander nucleotides are in black. Data are means ± SD from n=2 independent experiments. (E) Histograms reporting the editing efficiency obtained for the insertion of RM combinations in HBE cells homozygous for the F508del-*CFTR* gene. Cells were electroporated with SpRY-CBE4max mRNA and two sgRNAs. Editing was quantified by EditR tool^61^ on Sanger sequencing chromatograms. Data are means ± SD from n=2 independent experiments. (F) Deep sequencing analysis of off targets (OTs) identified through GUIDE-seq analysis. Cumulative A>G or C>T base conversions inside the sgRNA spacer are reported. Data are means ± SD from n=2 independent experiments.

To assess the editing specificity, as reported before^45,47–49^, we first performed a whole off-target genome analysis (GUIDE-seq^50^) with SpG or SpRY nucleases, followed by deep sequencing analysis of the nominated off-targets in cells treated with base editors. A limited number of off-target (OT) sites were revealed by SpG targeting the I539T locus (n=5), or SpRY targeting R555K (n=1) and R1070W (n=9) loci. In contrast, a higher number of OTs (n=20) was retrieved with the sgRNA R553Q+6 (**Figure S5).** The deep sequencing analyses to verify base editing on the retrieved OTs was performed in primary HBE cells by examining OTs with at least ≥30 sequencing reads in GUIDE-seq, thus largely over background levels. The percentage of base conversion reported in **Figure 5F** is the sum of all adenines or cytosines modified into the sgRNA spacer sequence. We measured 10% cumulative A>G conversion at OT1 with SpG-ABEmax I539T+7, corresponding to intron 1 of *KCNT2* gene. SpRY-BE4max R553Q+6 showed three main off target sites, OT1, OT2 and OT4, among which the strongest was OT1, with a C>T conversion higher than the on target site, corresponding to intron 3 of the *GALNT6* gene. Finally, SpRY-BE4max R1070W+6 exhibited almost 8% C>T conversion in OT2 corresponding to intron 1 of the *FLRT2* gene. No indels were detected beyond the reported A>G or C>T conversions (**Figure S6**).

### Functional rescue of CFTR anion transport

Based on the editing efficiency and precision combined with the capacity of the RMs to rescue CFTR activity, we selected the R555K-R1070W RMs for further characterization to restore the F508del function. For this purpose, we analyzed terminally-differentiated pseudostratified epithelia generated from HBE cells under air-liquid interface (ALI) condition. To evaluate CFTR-dependent transepithelial ion transport, we measured short-circuit current (Isc) in a Ussing chamber-like system^51,52^.

In the first set of experiments, we blocked the sodium current mediated by ENaC with amiloride and then stimulated CFTR activity with the membrane-permeable analog CPT-cAMP. Subsequently, we added the selective CFTR blocker CFTR_inh_-172 to abolish CFTR current. CFTR-dependent chloride secretion, as indicated by the amplitude of the current elicited by CPT-cAMP (cAMP) and the extent of current drop resulting from CFTR_inh_-172 (I-172) addition, was larger in cells edited with the R1070W and R555K-R1070W compared to HBE cultures not-treated (NT) or treated with a scramble sgRNA (CTR) (**Figure 6A**). In particular, the drop caused by the inhibitor (DI-172) showed a 3 to 4-fold increase in CFTR function with respect to NT or CTR in both R1070W and R555K-R1070W edited cells. Conversely no significant enhancement could be observed with the single R555K.

**Figure 6.**
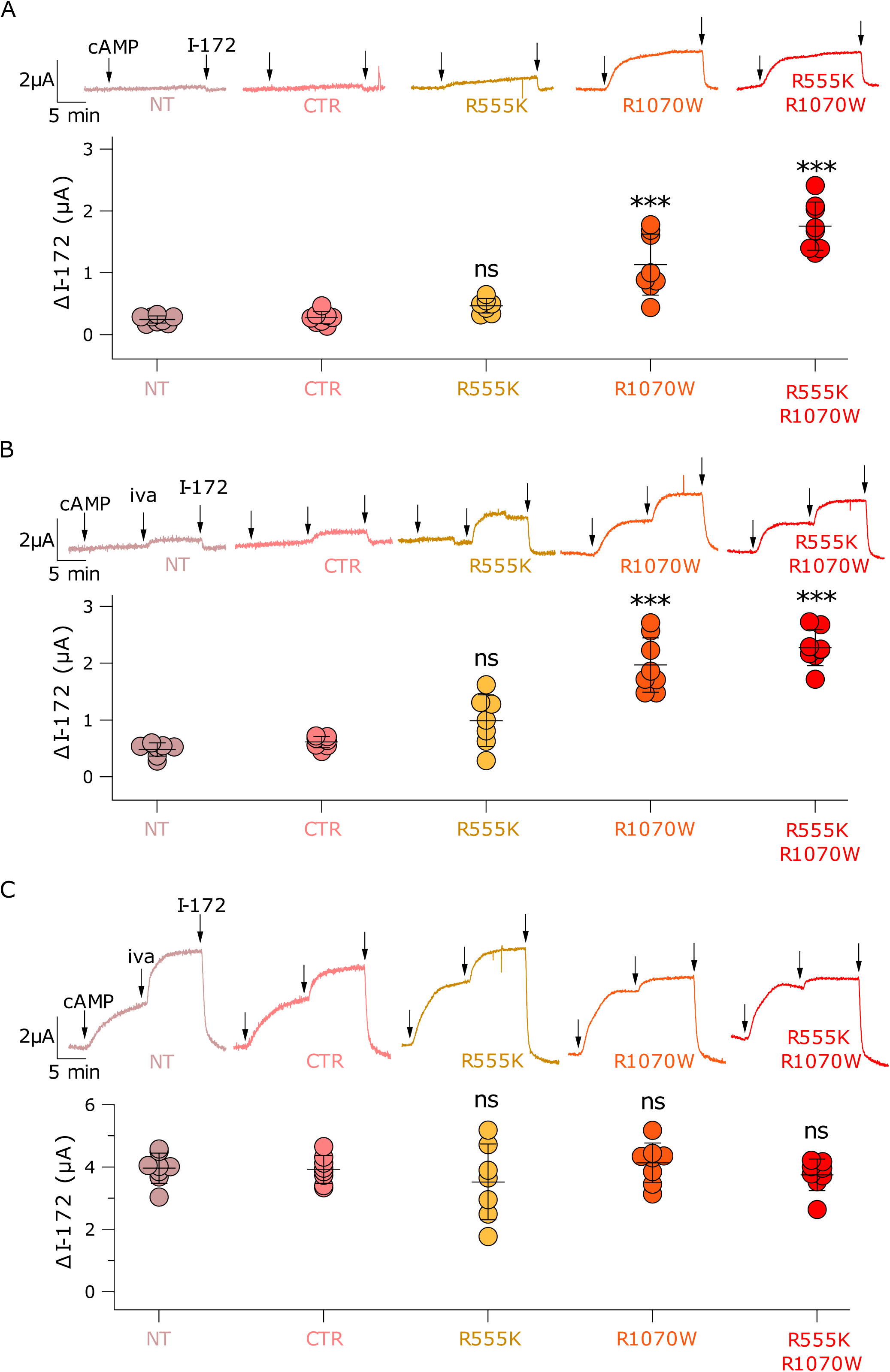
RMs insertion in HBE cells promote CFTR activity rescue. (A-C) Representative short-circuit current traces and summary of results obtained from epithelia generated from untreated cells (NT) or cells treated with SpRY-CBE4max-CTR or SpRY-CBE4max-sgRNAs. Editing efficiency is shown in **Figure S7**. For the representative traces, the identity of the compound addition (arrows) is shown for simplicity only in the far left trace and the initial part of the recording, containing the effect of amiloride, has been removed. In each panel, the scatter dot plots report the amplitude of the current drop elicited by CFTRinh-172 (I-172). (A) Recordings with sequential addition of CPT-cAMP (cAMP) followed by CFTRinh-172 (I-172). (B) Recordings in which 1 µM ivacaftor (iva) was added in the apical solution after CPT-cAMP. (C) Recordings in which epithelia were previously incubated with tezacaftor and elexacaftor for 24 h. Scatter dot plots represent the results from epithelia of n ≥ 7 of two independent experiments. Data are presented as means ± SD.

We also verified the potential additive effect that could be generated between RMs and ivacaftor, the potentiator molecule composing the ETI treatment. Ivacaftor, added after CPT-cAMP during Isc recordings, resulted in a further increase of CFTR-dependent current with a consequent enlargement of the effect of CFTR_inh_-172 (**Figure 6B**). The effect of ivacaftor was particularly evident in cells edited with R1070W and R555K-R1070W. An increase in current by ivacaftor was also evident in traces from cells edited with R555K. We calculated DI-172 for all conditions. In this set of experiments, cells edited with R1070W and R555K-R1070W again showed an improvement in DI-172 value (3-4 fold increase compared to controls). There was also a positive trend for R555K cells but this was not statistically significant (**Figure 6B**).

Finally, a third set of experiments was done with complete ETI by combining ivacaftor with the two correctors, tezacaftor and elexacaftor. Under this condition, CFTR function was similarly rescued irrespective of presence/absence of gene editing (**Figure 6C**). The lack of additive effects between pharmacological treatment and gene editing is likely due to the maximum F508del-CFTR rescue already obtained with ETI.

Overall, our data demonstrate that the insertion of RMs, specifically R1070W, in the F508del-*CFTR* locus of primary HBE cells efficiently rescues the CFTR ion-channel activity.

Finally, we also calculated the fraction of the total current that is activated by CPT-cAMP alone from the experiments performed in **Figure 6B** and **C**. The results, shown in **Figure 7A**, indicate that insertion of R1070W, significantly increases this parameter compared to R555K. These results suggest that R1070W, besides improving the trafficking of mutant CFTR to plasma membrane, also improves the gating as indicated by the proportionally reduced effect of ivacaftor.

**Figure 7.**
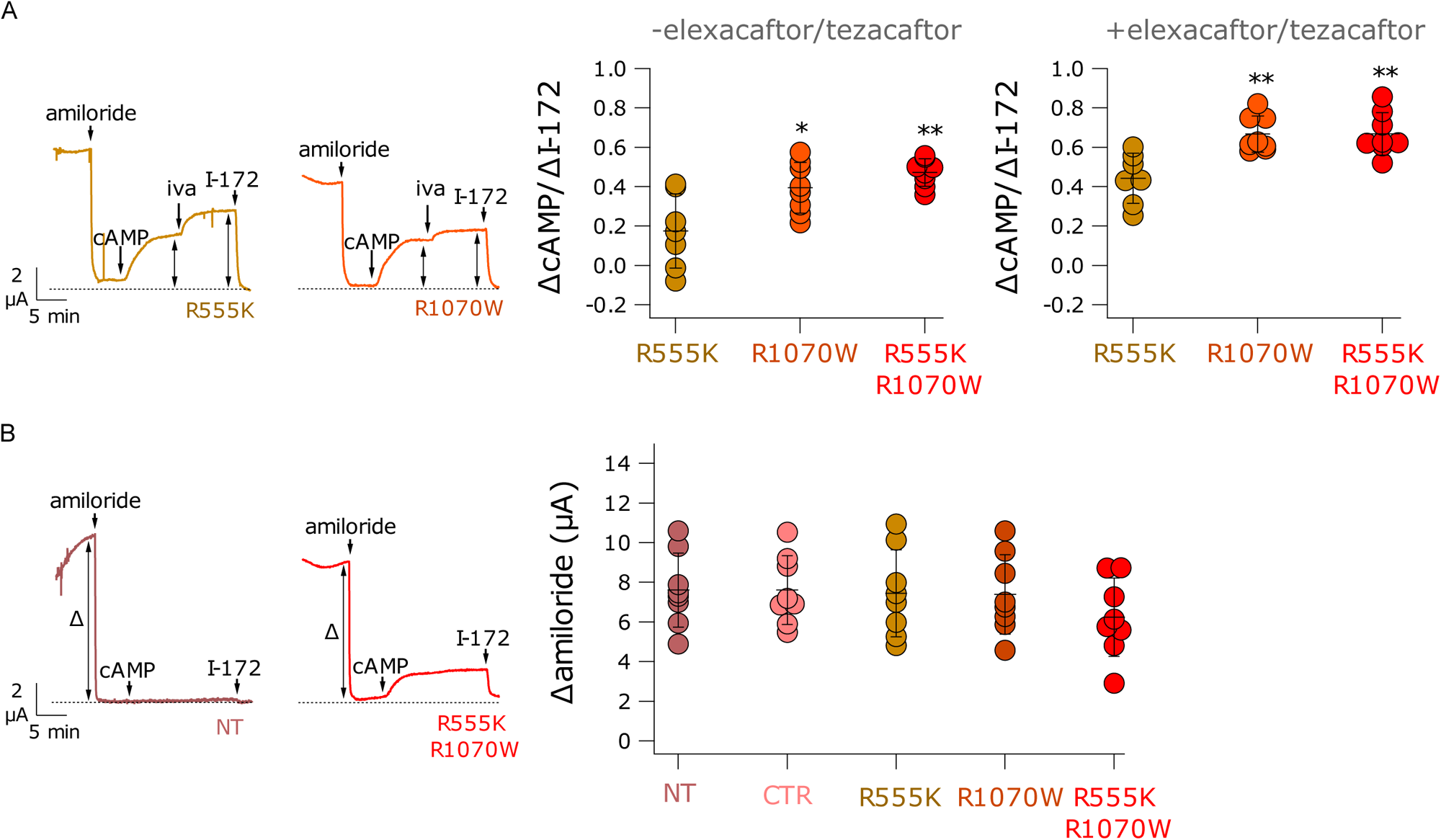
Analysis of gene editing effects on CFTR and ENaC properties. (A) Representative traces (left) and summary graphs (middle and right) showing the effect of gene editing on the relative response to CPT-cAMP (cAMP) and ivacaftor (iva). The scatter dot plots report the ratio between the current activated by CPT-cAMP alone and the total CFTR current activated by CPT-cAMP plus ivacaftor and blocked by CFTRinh-172 (I-172). Cells were previously treated with DMSO (middle) or with elexacaftor/tezacaftor (right). *, p < 0.05; **, p < 0.01 vs. R555K (ANOVA with Dunnett’s post hoc test). (B) Representative traces (left) and summary graph showing the activity of the ENaC channel. The scatter dot plot reports the amplitude of amiloride effect for the indicated conditions. No significant change in ENaC activity was observed by electroporation or by insertion of R555K and/or R1070W mutations. Data are presented as means ± SD.

We also evaluated the effect of gene editing on the effect of amiloride, added at the beginning of recordings. The amplitude of amiloride does not only reflect the activity of the sodium channel ENaC, but also in general the driving forces that underlie transepithelial ion transport. As shown in **Figure 7B**, the amplitude of the current blocked by amiloride was not significantly affected by the electroporation procedure (CTR) or by gene editing.

## Discussion

Technological advancements in genome editing offer new opportunities for treating genetic diseases like CF. Several strategies have been developed to repair CF causing mutations including the F508del^9^, the most frequent in CF patients^1^. In several studies the HDR capacity to substitute genomic fragments enhanced by nucleases was exploited to repair CF mutations including the F508del ^10–15^. Nonetheless, despite relevant advancements, HDR still remains a complex approach as it requires the delivery of a donor template and it is limited to dividing cells^53^. This poses significant obstacles for the treatment of the lung epithelium and potentially other tissues composed of differentiated and undifferentiated cells. Lungs are prioritized targets in CF as this is the organ mainly hampered by the disease. Nonetheless, the role of different cell types in the lung epithelium has not been defined yet, thus eluding the best cellular target for gene correction. Epithelial stem cells, while promising targets due to their fundamental role in tissue regeneration, present risks since nuclease activity associated with HDR and other genome editing strategies may negatively impact their staminal properties with potential downstream negative effects in derived tissues^54–56^. Moreover, DSBs generated by nucleases in HDR and other approaches have been reported to induce genomic rearrangements including translocation and chromosome aberrations^16–18^. In recent years more DSB free tools have been developed. Prime editing is an efficient genome editing strategy for both dividing and non-dividing cells and allows the introduction of all 12 possible types of point mutations, small insertions and small deletions^19^. This tool is composed of various elements requiring fine tuning for efficient and precise editing. A first attempt to correct the F508del locus through prime editing showed an overall low efficiency and high levels of indels introduced on both alleles due to the additional nicking sgRNA^20^. More recently an intensive optimization study produced an enhanced prime editing strategy to repair the F508del mutation^20,21^. The optimization comprises several modifications of the editor, including the co-expression of a non physiological dominant-negative mismatch repair protein (MLH1dn), which resulted in recovery of CFTR activity similar to levels obtained with pharmacological treatments. The correction of the mutation was associated with indels that, even though less than previous prime editing or HDR approach, still are remarkably high (around 10% at highest editing conditions).

Base editing has emerged as another promising strategy for correcting CF mutations^45,57–59^. It is usually more efficient than prime editing^20,21^ and is better suited for delivery due to reduced complexity compared to HDR, requiring a co-delivery with donor DNA template, and prime editor, using two guide RNAs (pegRNA and ngRNA) plus a cofactor (MLH1dn) to enhance the editing.

Nonetheless, base editors are limited to A>G or C>T transitions, making it unsuitable for correcting deletions like the F508del. To overcome this limitation, we explored the use of RMs to compensate for the F508del-CFTR defect^24^. We selected RMs compatible with a base editing strategy, to be introduced into the F508del allele as single or combined mutations. I539T, G550E, R553Q, R555K, located in the NBD1 domain, are mainly responsible in restoring the protein folding^26–31^, while the R1070W, at the TMD2 domain, increases PM levels and also improves protein gating^24,30,32^. The combination of R555K and R1070W mutations proved particularly effective, as they act at different stages of CFTR biogenesis and complement each other in restoring CFTR function. The success of this approach was further enhanced by an opportunistic bystander edit that introduced an additional beneficial mutation, R553Q^60^, resulting in a triple modification (R553Q-R555K-R1070W) using a single base editor and two sgRNAs.

A second bystander edit associated with this RMs combination was observed with the sgRNA for R1070W producing a silent mutation thus very likely innocuous. Nonetheless, the bystander edits were lowered to background levels in primary HBE cells, probably as a result of transient expression of base editors delivered as RNA, as reported before^45^. The off-target profile for selected sgRNAs was quite heterogeneous with diverse percentages of non-specific modifications. Focusing on the R555K-R1070W RMs we detected exclusively a non-specific edit in the intronic region of the *FLRT2* gene which would deserve further investigation to determine the impact in the target cells. The deep sequence analysis showed no indels formation at the target site thus precluding genotoxic activity as observed for HDR and prime editing.

The capacity of the R555K-R1070W, introduced through base editing, to recover F508del-CFTR function was validated in terminally-differentiated HBE cells derived from patients by measuring the short-circuit current in pseudostratified epithelia grown in culture. The genome editing strategy alone recovered CFTR activity, while not altering the sodium channel ENaC, as well as more general transepithelial ion transport, as observed by the response to the amiloride. A synergistic effect between the editing and pharmacological treatment was observed by using potentiators, molecules improving the CFTR gating, in particular with the R555K-R1070W or R1070W alone. These results indicate that the recovery of F508del-CFTR plasma membrane proximity by editing is sufficient to retrieve normal current flows by the potentiator treatment. Conversely, pharmacological cocktails containing both correctors and potentiators (ETI) reached the maximum enhancement of F508del-CFTR recovery, and no benefits could be generated by the RMs. Nonetheless, CFTR activity analyzed in cell lines expressing the F508del-CFTR modified with RMs showed an incremental effect of both correctors and potentiators with the majority of the editing strategies. These results suggest that increasing the editing efficacy through more advanced tools may lead to powerful combinatorial treatments increasing the therapeutic efficacy of the CFTR modulators to lower the drug dosage regimen.

Even though CFTR modulators have significantly improved the patients’ quality of life and extended the lifespan, the burden of lifelong and costly treatments, which are also occasionally inefficacious, highlight the urgent need for a more definitive cure. Indeed, with this study we first prove that beyond the value of potential permanent cure for CF, genome editing offers the opportunity to design a combinatorial treatment to increase the treatment solutions.

Harnessing the CFTR folding and activity by targeting specific domains is a new frontier for the treatment of CF.

## Acknowledgements

We are grateful to Cereseto’s lab members for helpful discussion throughout the project. We thank the Next Generation Sequencing facility at the University of Trento for technical support.

We are grateful to Prof. Marianne Carlon for sharing reagents and for precious insights in CFTR testing.

This work was fully supported by the Italian Cystic Fibrosis Foundation for Research (FFC) through the FFC#3-2019, FFC#2-2021 and GenDel-CF.

## Author contributions

I.C., G.M., C.G, A.U, D.G., M.D.S. designed and performed the experiments; I.C., G.M., C.G, A.U, D.G., M.D.S., L.J.V.G. and M.C. collected and analyzed the data; G.P., G.M., D.A. and A.C. conceived and designed the study, I.C., G.M., A.U, L.J.V.G., D.G., D.A. and A.C. wrote and edited the paper; A.C. and D.A. were responsible for the coordination of the study. All authors read, corrected, and approved the final manuscript.

## Declaration of interests

A.C. and G.P. are co-founders and hold stocks of Alia Therapeutics, a genome editing company.

## Methods

### Plasmids

pCHMWS-CFTR-wt-ires-PURO and pCHMWS-CFTR-F508del-ires-PURO plasmids (kindly provided by Prof. Marianne Carlon, KU Leuven Belgium) contain a triple hemagglutinin (3HA) tag in the fourth extracellular loop of CFTR and a puromycin resistance.

pLenti-CFTR-wt-ires-PURO and pLenti-CFTR-F508del-ires-PURO plasmids were cloned starting from a pLenti-FNLS-P2A-Puro plasmid (Addgene, #110841), substituting the BE4max-P2A-Puro sequence with CFTR-ires-PURO derived by PCR on the pCHMWS-CFTR plasmids, using BamHI and MluI restriction enzyme digestion.

F508del-CFTR RMs were cloned into the pLenti-CFTR-F508del-ires-PURO plasmid using PCR amplification and BsmBI restriction enzyme. Oligonucleotides used for cloning are listed in **Table S1**.

Plasmids expressing sgRNAs were cloned in a pUC19-sgRNAopt (pUC19 plasmid with sgRNA cassette derived from pX330, Addgene, #42230) as described before^62^. SgRNA spacers are reported in Table S2.

HS-YFP (YFP-H148Q/I152L/F46L) plasmid encode for halide-sensitive YFP was used for YFP assay.

In base editing experiments the following plasmids were cloned by restriction enzyme strategies: pCMV-SpG-ABEmax, pCMV-SpG-ABE8e, pCMV-SpG-ABE8.20m, pCMV-NG-ABEmax^45^, pCMV-NG-ABE8e^45^, pCMV-AncBE4max-SpG, pCAG-CBE4max-NG-P2A-EGFP, pCMV-AncBE4max-NG. pCAG-CBE4max-SpRY-P2A-EGFP (#139999) and pCAG-CBE4max-SpG-P2A-EGFP (#139998) were purchased from Addgene.

pIVT-CMV-SpG-ABEmaxP2A-EGFP and pIVT-CBE4max-SpRY-P2A-EGFP, used for in vitro RNA transcription, were constructed starting from pCMV-SpG-ABEmax and pCAG-CBE4max-SpRY-P2A-EGFP introducing the ABE/CBE-Cas9 sequence in the pIVT-SpRY-ABE9 plasmid by EcoRI restriction enzyme digestion. Oligonucleotides used for cloning are listed in Table S3.

For GUIDE-seq analyses pCMV-T7-SpG-P2A-EGFP (Addgene #139998) and pCMV-T7-SpRY-P2A-EGFP (Addgene #139989) plasmids were used.

### Cell lines

HEK293T cells were obtained from the American Type Culture Collection (ATCC; www.atcc.org). HEK293T stably expressing HS-YFP, wtCFTR and F508del-CFTR were kindly provided by Prof. MS Carlon (KU Leuven Belgium). HEK293T cell lines stably expressing the different CFTR variants were generated by lentiviral transduction.

Cells were cultured in Dulbecco’s modified Eagle’s medium (DMEM; Life Technologies) supplemented with 10% fetal bovine serum (FBS; Life Technologies), 10 U/mL antibiotics (PenStrep; Life Technologies), and 2 mM L-glutamine at 37° C in a 5% CO2 humidified atmosphere.

### Lentiviral production and transduction of cell lines

Lentiviral particles were produced in HEK293T cells at 80% confluency in 100mm dishes. 10 µg of pLenti-CFTR-F508del-RMs-ires-PURO plasmid, 6.5 µg of Δ8.91 packaging plasmid and 3.5 µg of VSV-G were transfected using polyethylenimine (PEI). The viral supernatant was collected after 48 h and filtered in 0.45 μm PES filter and aliquots were stored at −80°C. Vector titers were measured as reverse transcriptase units (RTU) by SG-PERT method64.

For transduction experiments, 150.000 HEK293T cells were seeded in a 24-well plate and transduced 24 after hours with 1 RTU of lentiviral vectors. 48 h later, cells were selected with puromycin (2 μg/ml). Cells were kept under puromycin selection and experimental analyses were performed 3 weeks after transduction.

### Cell lines transfection

HEK293T cells or HEK293T stably expressing HS-YFP, were co-transfected with 750 ng of base editor and 250 ng of sgRNA encoding plasmids. Transfections were performed using Transit LT1 reagent (Mirus Bio) according to the manufacturer’s protocol. Cell pellets were collected 3 days post-transfection for editing analyses.

### Editing analyses

Genomic DNA was extracted using QuickExtract DNA extraction solution (Epicentre). Target regions were amplified by PCR with HOT FIREPol® DNA Polymerase (Solis BioDyne) using 100–500 ng of DNA as template. Oligos are listed in Table S4, S5. PCR products were purified using NucleoSpin Gel and PCR Clean-up (Macherey-Nagel) and sequenced by Sanger sequencing. To evaluate editing efficiency, the chromatograms were analyzed using EditR software^61^.

### Western Blotting

Cell pellets were collected from cells stably expressing CFTR variants and lysed with RIPA buffer. 20 μg of the whole lysates quantified by BCA assay (Thermo Fisher Scientific) were separated by SDS-PAGE in a 6% polyacrylamide gel and then transferred to a PVDF membrane. Membranes were probed with a mix of 4 mouse monoclonal CFTR antibodies purchased from the CFF Foundation (Ab 570 and 596 diluted 1:500; Ab 660, TJA9 diluted 1:1000) and mouse anti-α-actinin (1:1000, sc-17829, Santa Cruz). Detection was performed with Pierce ECL reagent (Thermo Fisher Scientific) using the Uvitec Alliance Instrument.

### Flow Cytometry

Cells were stained with HA.11 antibody (#901515, Biolegend 1:1000) at 4°C for 45 minutes, followed by Alexa-488 secondary antibody (#A-11001, Thermo Fischer Scientific, 1:500) for 30 minutes and then analyzed at BD FACS Canto™, as previously described^63^. Cells not expressing CFTR were used as a negative control to determine the fraction of positive cells. WtCFTR and F508del-CFTR cells were used to determine the fraction of cells expressing CFTR at the PM (**Figure S1**).

### Halide Sensitive (HS) YFP Assay

All transfections were performed using Transit LT1 reagent (Mirus Bio) according to the manufacturer’s protocol. HEK293T stably expressing HS-YFP were transfected with 750 ng of plasmids encoding CFTR wt, F508del or F508del-RMs. 24 hours after transfection, cells were seeded into 96-well plates pre-coated with Poly-D-Lysine (#A3890401, ThermoFisher). After 24 hours, HS-YFP quenching assays were conducted as follows.

Cells were washed with DPBS (NaCl 137mM, KCl 2.7mM, Ca2Cl 0.7mM, MgCl2 1.1mM, KH2PO4 1.5mM, Na2HPO4 8.1mM) and incubated in 60 µl of DPBS containing either the CFTR activator forskolin (#S2449 Selleckchem, 10 µM) or DMSO as a negative control for 20 minutes at 37°C. Fluorescence was measured using an inverted epifluorescence microscope equipped with a 490 nm excitation filter (Chroma Technology, ET490/20x) and a 540 nm emission filter. Continuous fluorescence recording was performed for 35 seconds. After an initial 5-second recording, 165 µl of I− buffer (NaI 137mM, KCl 2.7mM, CaCl 0.7mM, MgCl2 1.1mM, KH2PO4 1.5mM, Na2PO4 8.1mM) was added fluorescence was monitored for an additional 30 seconds. YFP quenching was calculated as the ratio F/F0 at the end of the recording interval and CFTR function was expressed as 1−(F/F0).

For analysis of CFTR recovery induced by base editing, 150.000 HEK293T cells stably expressing HS-YFP were seeded into 24-well plates and transfected with 750 ng of base editor and 250 ng of sgRNA. 24 hours post-transfection,cells were moved into 96-well plates, and HS-YFP was performed after an additional 24 hours.

To evaluate the synergistic effects between revertants and pharmacological modulators, HEK293T cells stably expressing CFTR variants were transfected with 250 ng of plasmid encoding HS-YFP. The following day, cells were seeded in 96-well plates and incubated with either lumacaftor (#S1565, Selleckchem, 2.5 µM) or DMSO for 24 hours. After overnight incubation, cells were washed with DPBS, and a solution containing both the potentiator ivacaftor (#S1144, Selleckchem, 3 µM) and forskolin (10 µM) were added for 20 min before starting the HS-YFP quenching assay.

### *In vitro* mRNA transcription

SpG-ABEmax and SpRY-BE4max mRNAs were synthesized by T7 polymerase *in vitro* transcription using HighYield T7 RNA Synthesis Kit (Jena Bioscience). In detail, 750 ng of linearized pIVT-SpG-ABEmax-EGFP and pIVT-SpRY-BE4max-EGFP plasmids were used as template for the reaction. CleanCap Reagent M6 (TriLink BioTechnologies, 15mM) was added to the reaction. mRNA was purified with LiCl precipitation and resuspended in DEPC-ddH2O. The sgRNAs (standard or with chemical modifications^62^) were synthetically ordered and produced from Integrated DNA technologies (IDT) Table S6.

### Electroporation of cell lines

200,000 HEK293T cells stably expressing F508del-CFTR were electroporated with 3 μg of base editor mRNA either alone (Ctr) or with 3 μg of sgRNA. Electroporation was performed with the Amaxa 4D-Nucleofector (SE Cell Line buffer, CM-130 program). Next cells were seeded into a 12-well plate and collected after 3 days for editing analysis.

### Primary bronchial epithelial cell culture and electroporation

Primary bronchial epithelial cells (HBE) were derived from a CF patient homozygous for the F508del and were provided by the Primary Cell Culture Service of the Italian Cystic Fibrosis Research Foundation. The Ethics Committee of the Istituto Giannina Gaslini approved this study and informed consent was obtained from all participating CF subjects. Cells were cultured in LHC9/RPMI 1640 (1:1) without serum as previously described^51,52^ supplemented with rho-associated protein kinase 1 inhibitor (Y-27632, Merck, 5 mM), SMAD-signaling inhibitors, bone morphogenetic protein antagonist (DMH-1, Merck, 1 mM), and transforming growth factor β antagonist (A 83-01, Merck, 1 mM)^64^. 200.000 cells were electroporated with the Amaxa 4D-Nucleofector (P3 primary cells buffer, DN-100 program) and seeded into a 12-well plate previously treated with collagen.

Several amounts of base editor mRNA and sgRNA R553Q+6 (standard or with chemical modifications^62^) were tested in electroporation experiments to find the most efficient condition (Figure S7). After the initial setting, the electroporation was performed for all the other revertants using 6 μg of base editor mRNA either alone or with 6 μg of sgRNA scramble (CTR) and sgRNAs to insert I539T, R553Q, R555K, R1070W. After 3 days cells were collected for editing analysis.

### Short-circuit current recordings

HBE cells, resuspended in LHC9/RPMI 1640 proliferative medium, were seeded at high density on Transwell (cc3470, Corning Costar; 0.33 cm^2^ surface) porous inserts (150.000 cells per insert). After 24 h, to achieve the air-liquid interface condition, the medium on the apical side was totally removed, whereas the medium on the basolateral side was replaced with PneumaCult ALI (Stemcell Technologies) differentiation medium. Short-circuit current experiments were performed after 2–3 weeks in culture, when epithelia were fully differentiated. Transwell supports carrying the epithelia were mounted in vertical chambers with internal fluid circulation (EM-CSYS-8, Physiologic Instruments). The apical and basolateral compartments were filled with a solution containing in mM: 126 NaCl, 0.38 KH2PO4, 2.13 K2HPO4, 1 CaCl2, 1 MgSO4, 24 NaHCO3, 10 glucose, and phenol red. Solutions were continuously bubbled with a mixture of 5% CO2-95% air and kept at 37 °C. The transepithelial voltage was clamped at 0 mV with an eight-channel voltage-clamp amplifier (VCC MC8, Physiologic Instruments, San Diego, CA, USA) connected to apical and basolateral compartments with Ag/AgCl electrodes through agar bridges (1 M KCl in 2% agar). The resulting short-circuit current was recorded with the Acquire & Analize 2.3 software (Physiologic Instruments, San Diego, CA, USA). During experiments, epithelia were sequentially treated with amiloride (10 µM, apical) to block ENaC-dependent current, CPT-cAMP (100 µM, apical and basolateral) to stimulate CFTR activity, and CFTRinh-172 (20 µM, apical) for CFTR inhibition.

### GUIDE-seq

GUIDE-seq analysis was performed as previously described^45,65^. In short, HEK293 cells were transfected with 500 ng of SpGCas9 or SpRYCas9 plasmids, 250 ng of sgRNA plasmids, 10 pmol of dsODNs and 50 ng pEGFP-IRES-puro plasmid using Lipofectamine 3000 (Invitrogen^TM^). 24h after transfection, cells were split and selected with 1 mg/ml puromycin. Genomic DNA was extracted (DNeasy Blood and Tissue kit, #69506, Qiagen) and sheard to average length of 500 bp using sonication (Focused-ultrasonicator, Covaris). End-repair and sequencing adaptor ligation were performed using NEBNext Ultra End Repair/dA Tailing Module (E7546S, New England Biolabs) and NEBNextUltra Ligation module (E7595S, New England Biolabs respectively following the procedures reported by Nobles et al.^66^. Amplification steps were then performed following the GUIDE-seq protocol from Tsai et al.^50^ and libraries were quantified using Qubit dsDNA High Sensitivity Assay kit (Q33233, Invitrogen^TM^). Next-generation sequencing was performed on an Illumina MiSeq sequencing device using Illumina Re-agent kit V2-150PE. Raw sequencing data (FASTQ files) were analyzed using the GUIDE-seq computational pipeline (v1.0.2) available at https://github.com/tsailabSJ/guideseq. GUIDE-seq data are available in Table S7. Raw sequencing data have been submitted to NCBI-SRA, BioProject accession number PRJNA1150519.

### Targeted deep sequencing

Genomic DNA was extracted from HBE cells (F508del/F508del) 3 days after electroporation. The loci of interest (on and off targets) were amplified using Phusion high-fidelity polymerase (Thermo Scientific). Amplicons were indexed by PCR using Nextera indexes (Illumina), pooled in equimolar concentrations, and sequenced on an Illumina Miseq system using an Illumina Miseq Reagent kit V-300 cycles (2 x 150 bp paired end). Primers used for PCR amplification are listed in Table S8. FASTQ data were analyzed using CRISPResso version 2.3.1^67^. Raw sequencing data have been submitted to NCBI-SRA, BioProject accession number PRJNA1150519.

### Statistical analysis

All statistical analyses were performed by GraphPad Prism Prism 10 software. For HBE cells ordinary one-way analysis of variance (ANOVA) was performed, using Dunnett’s post hoc test. Statistical significance was defined as * p<0.05, ** p<0.01, *** p<0.001, **** p<0.0001, and ns, not-significant.

## Supplemental Information

**Document S1. Figure S1-S7**.

**Table S1.** Oligonucleotides used to clone F508del-CFTR RMs into the pLenti-CFTR-F508del-ires-PURO plasmid.

**Table S2.** Sequence of sgRNA spacers.

**Table S3**. Oligonucleotides used for pIVT plasmids cloning.

**Table S4.** Oligonucleotides used to evaluate editing efficiency in F508del-*CFTR* HEK293T cells.

**Table S5**. Oligonucleotides used to evaluate editing efficiency in BE cells.

**Table S6.** List of sgRNA sequences ordered from IDT.

**Table S7.** Guide seq data.

**Table S8.** Oligonucleotides used for targeted deep sequencing.

