## Supplementary material for "Functional rescue of F508del-CFTR through revertant mutations introduced by CRISPR base editing": Supplemenatry Information

#### Supplementary Figure 1

A

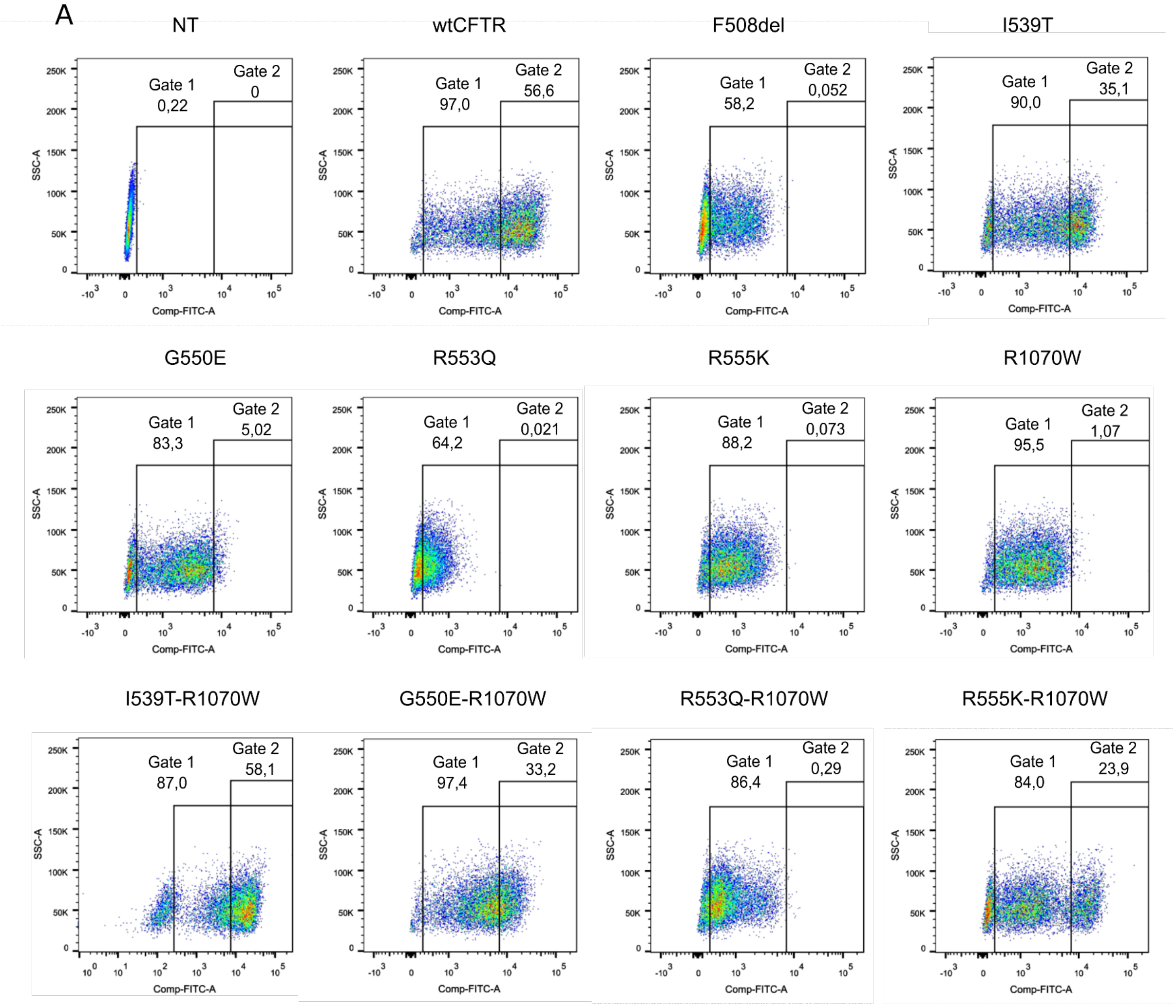


B

| REVERTANT | EXON | MUTATION | EDITOR |
|---|---|---|---|
| **I539T** | 12 | A>G | ABE |
| **G550E** | 12 | C>T | CBE |
| **R553Q** | 12 | C>T | CBE |
| **R555K** | 12 | C>T | CBE |
| **R1070W** | 20 | C>T | CBE |

**Figure S1.**
**(A)** Dot plots of FACS analysis of HEK293T cells transduced with RMs. HEK293T were stained with an anti HA primary antibody followed by an Alexa-488 secondary antibody. Cells not expressing CFTR were used as a negative control. F508del-CFTR and wtCFTR cells were used as reference to determine the fraction of cells expressing CFTR at the PM. As shown, F508del CFTR cells are mainly distributed in gate 1, while wtCFTR cells are mainly localized in gate 2. Revertant cell population localizes in gate 2. **(B)** Table of RMs inserted by base editors, including their location within the *CFTR*

gene, the specific base substitutions performed at the target site and the type of editor used for the insertion.

### Supplementary Figure 2

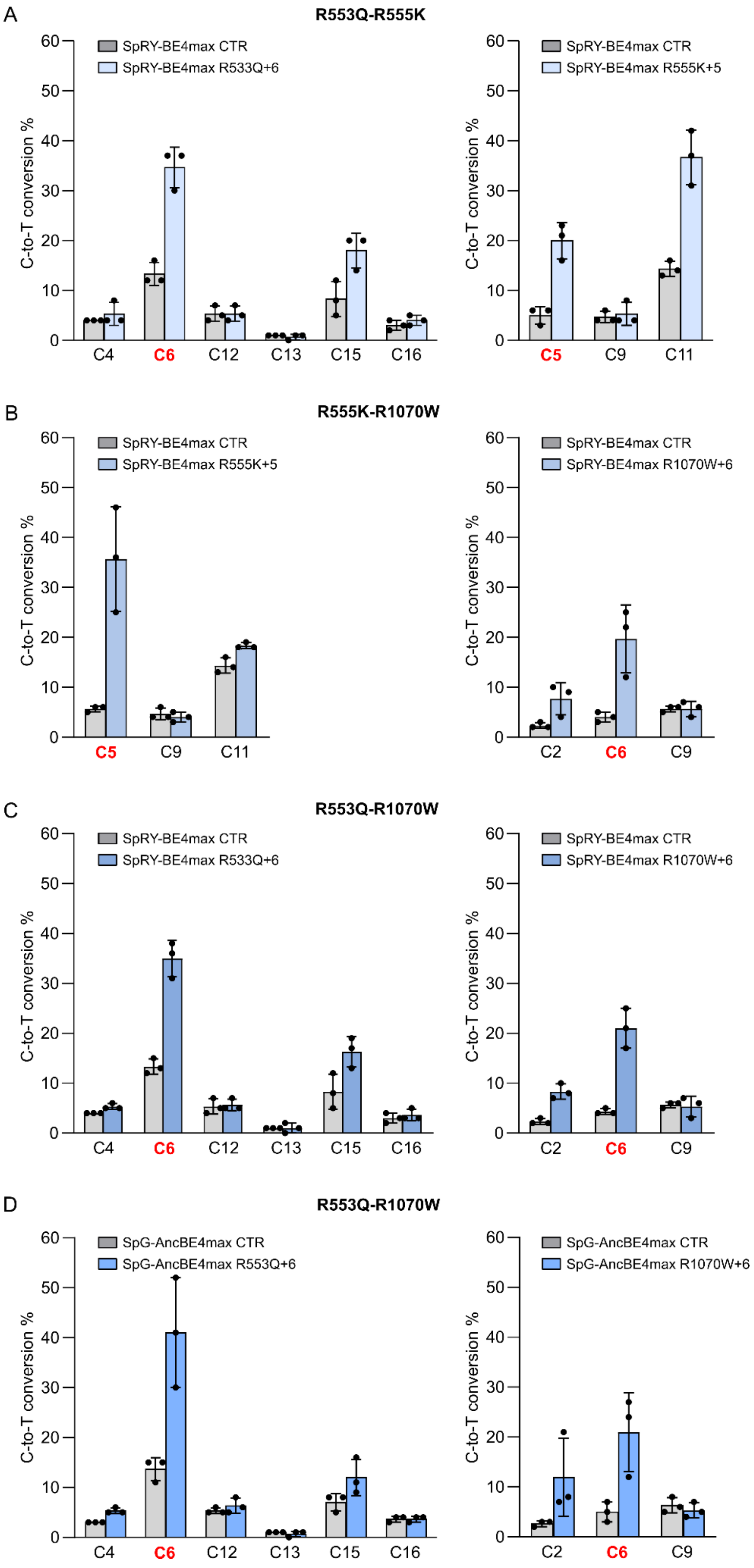

**Figure S2. (A-D)** Editing efficiency with bystander effect of the F508del-CFTR cDNA in HEK293T cells transfected with the indicated base editors and sgRNAs used in combination to generate **(A)** R553Q-R555K **(B)** R555K-R1070W **(C and D)** R553Q-R1070W RMs (as in Figure 2F). Editing was quantified by EditR tool after PCR amplification and Sanger sequencing of the target locus. The target base for each RM is highlighted in red; bystander edits are shown in black. Data are means ± SD from n=3 independent experiments.

### Supplementary Figure 3

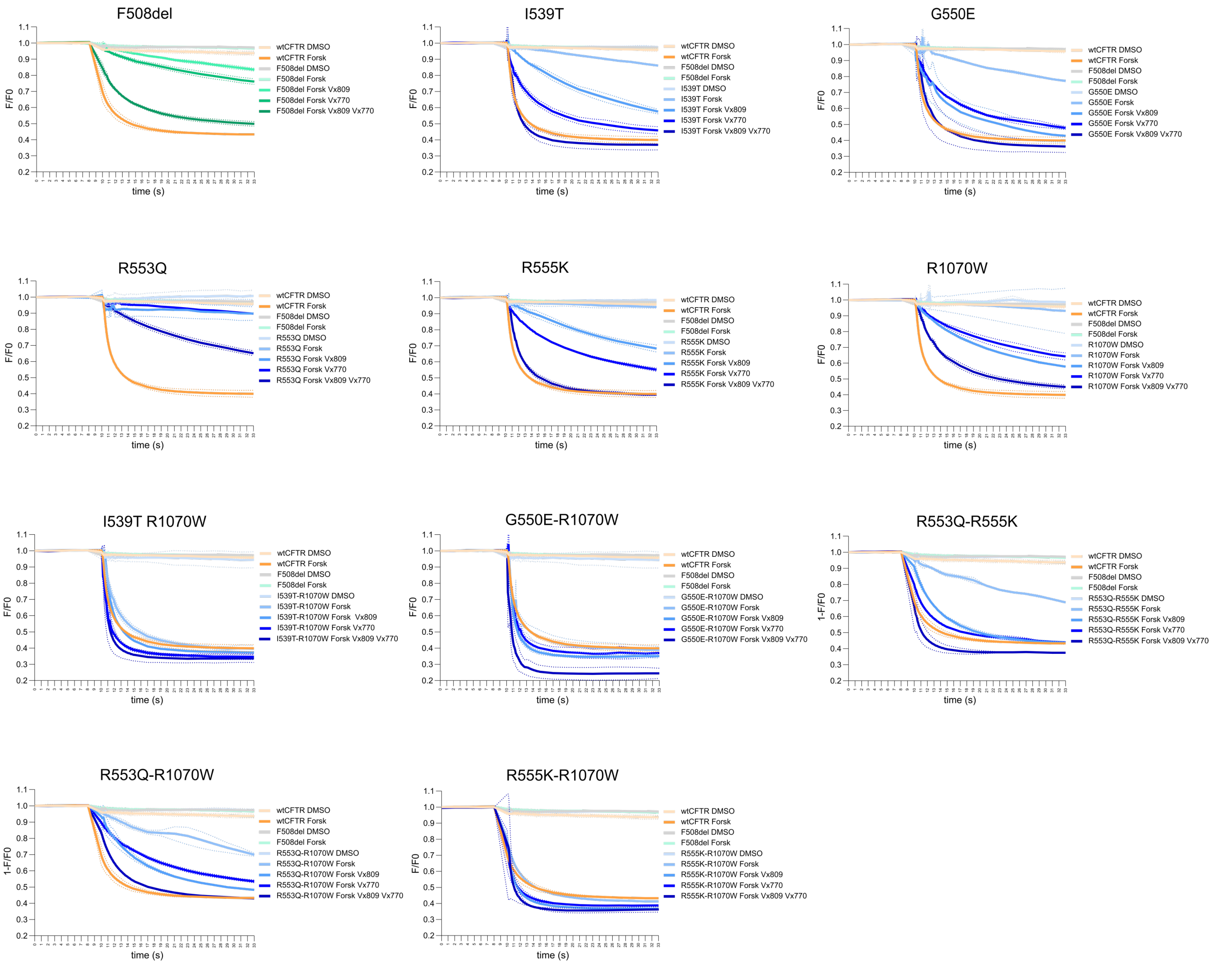


**Figure S3.** Representative traces of halide-sensitive yellow fluorescent protein (HS-YFP) quenching assay recorded with a fluorescence microscope. The assay was performed in HEK293T stably expressing F508del-CFTR variants and transfected with HS-YFP. The day after transfection cells were seeded in a 96-well plate and incubated with VX-809 (2.5 µM) or DMSO for 24 hours. After overnight incubation, cells were washed with DPBS, then potentiator VX-770 (3 µM) and forskolin (10 µM) were added for 20 min and HS-YFP quenching was performed. Graphs depict the mean of n=2 independent experiments.

#### Supplementary Figure 4

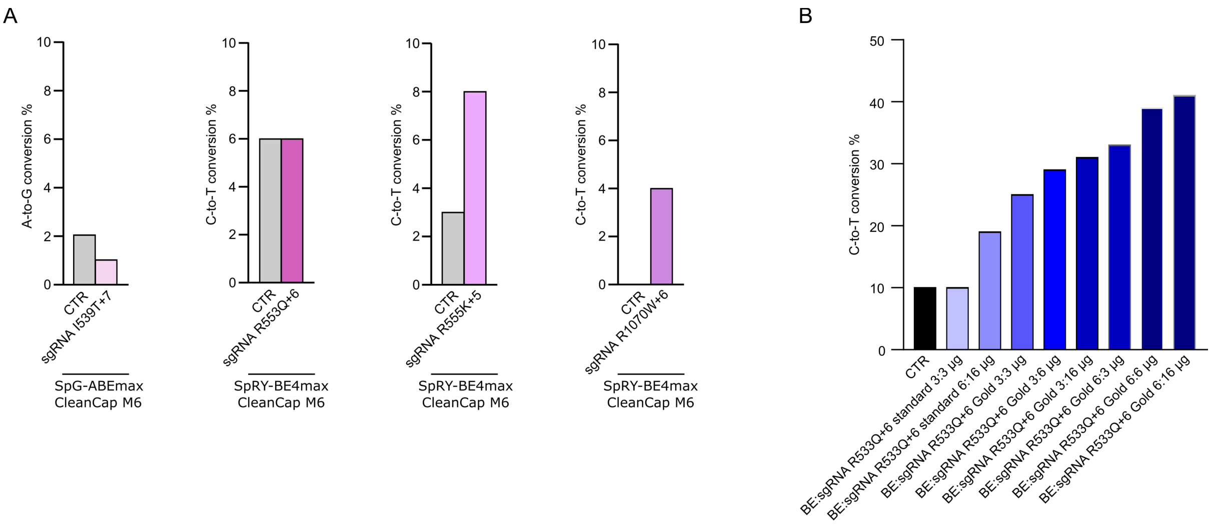


**Figure S4. (A)** Editing efficiency of RMs inserted into HBE cells homozygous for the F508del-CFTR gene. Editing was quantified by EditR tool of Sanger sequencing chromatograms. HBE cells were electroporated with 3 µg of ABE or CBE mRNA and 3 µg of standard sgRNA specific for the target base. **(B)** Enhanced editing efficiency following optimization of the experimental conditions. Chemical modifications were applied to sgRNAs and several BE/sgRNA ratios were tested to further refine the editing process.

#### Supplementary Figure 5

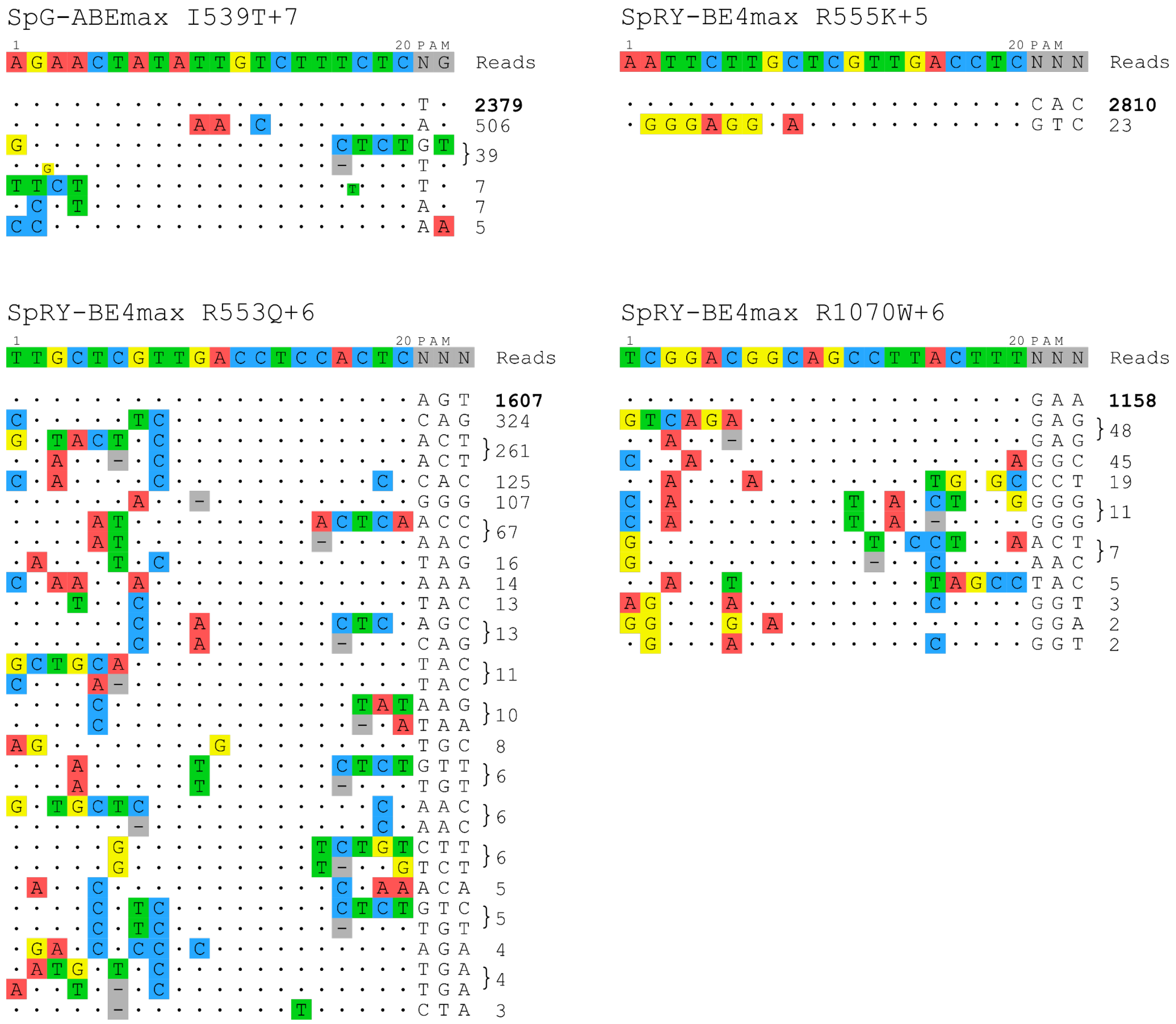


**Figure S5.** OT sites detected by GUIDE-seq with SpG and SpRY with sgRNAs targeting the genomic loci of interest. Each detected OT is represented with its corresponding number of GUIDE-seq reads. ON target reads are in bold.

### Supplementary Figure 6

I539T OFF Target 1 (Intron 1 of KCNT2 gene)

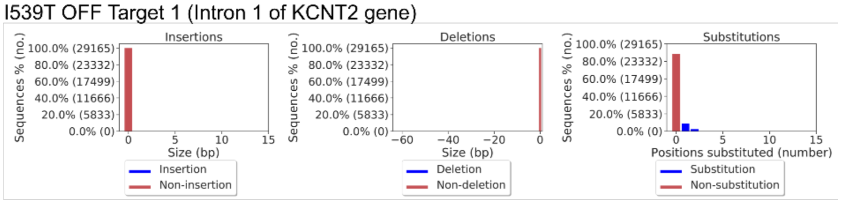


I539T OFF Target 2 (Intron 4 of ARID5B gene)

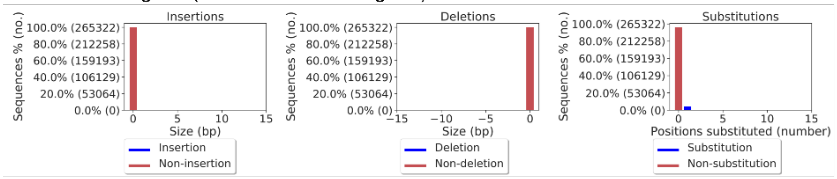


R553Q OFF Target 1 (Intron 3 of GALNT6 gene)

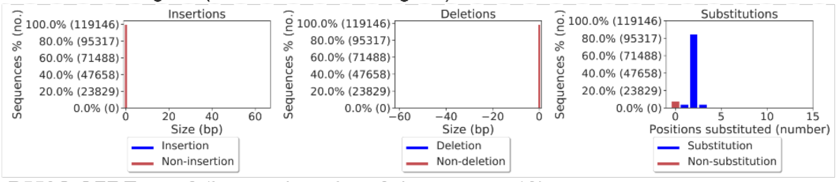


R553Q OFF Target 2 (Intergenic region of chromosome 10)

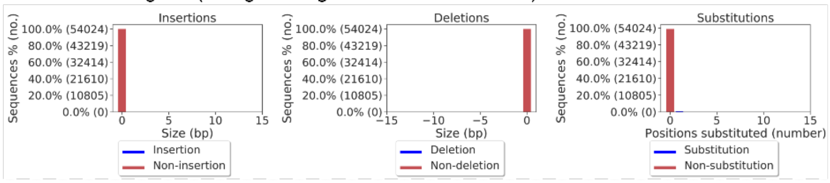


R553Q OFF Target 3 (Intron 4 of DNAJC5 gene)

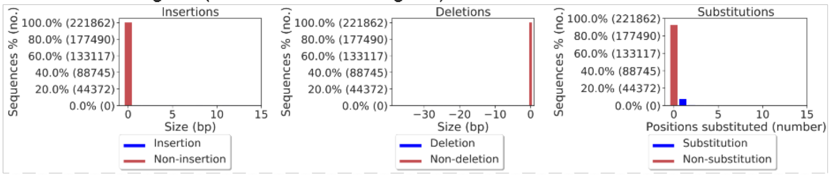


R553Q OFF Target 4 (Intron2 of LINC01164 gene )

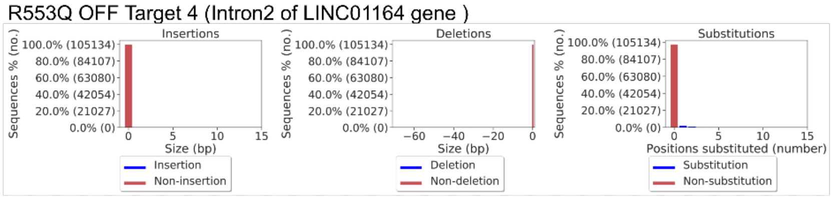


R553Q OFF Target 5 (Intergenic region of chromosome 9)

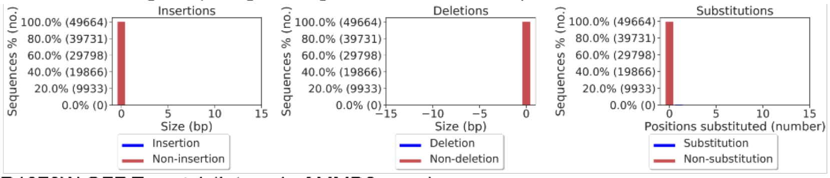


R1070W OFF Target 1 (Intron 1 of MMD2 gene)

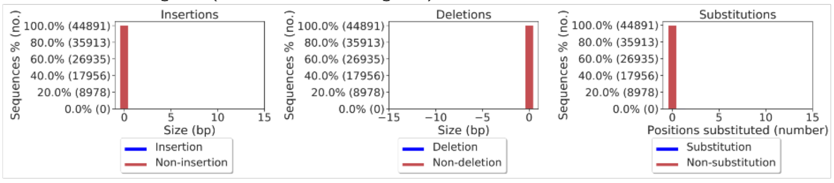


R1070W OFF Target 2 (Intron 1 of FLRT2 gene)

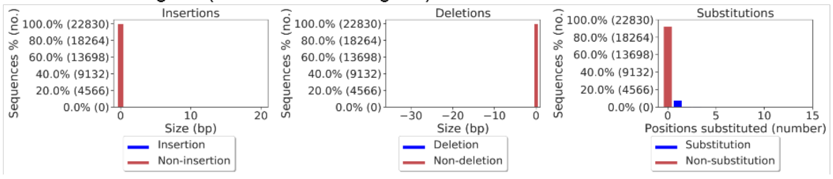

**Figure S6.** Deep sequencing analysis of OTs. Histograms report the percentage of insertions, deletions and base substitutions in each OTs as determined by deep sequencing analyses through CRISPResso.

#### Supplementary Figure 7

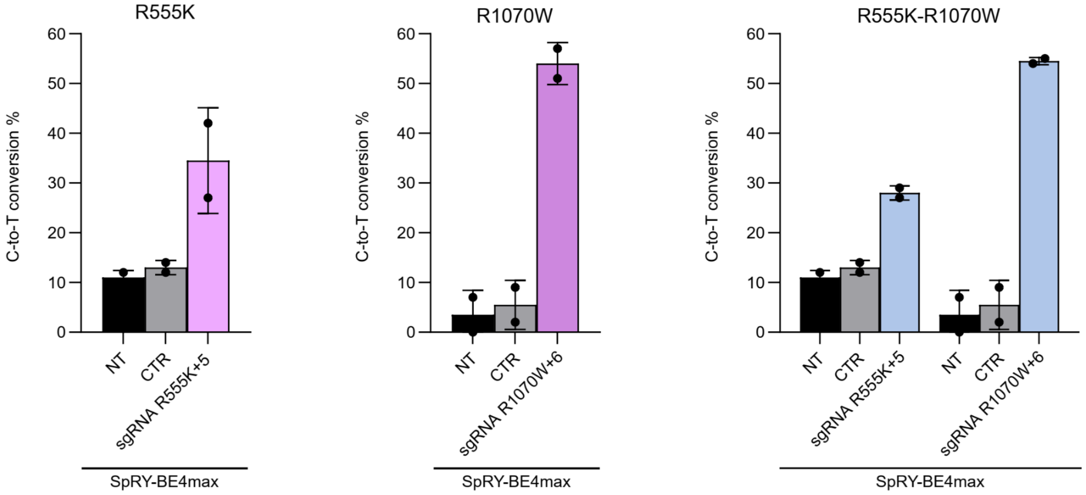


**Supplementary Figure 7.** Percentage of C-to-T conversion for the insertion of single and double RMs in HBE cells homozygous for the F508del-CFTR gene, used for CFTR function measurements (see Figure 6 and 7). HBE were electroporated with mRNA encoding the indicated base editors and chemical modified sgRNAs specific for the target nucleotide. Editing was quantified by EditR on Sanger sequencing chromatogram. Data are means ± SD from n=2 independent experiments.
